# Neutralization of a Distributed Coulombic Switch Precisely Tunes Reflectin Assembly

**DOI:** 10.1101/456442

**Authors:** Robert Levenson, Colton Bracken, Cristian Sharma, Jerome Santos, Claire Arata, Phillip Kohl, Youli Li, Daniel E. Morse

## Abstract

Reflectin proteins are widely distributed in reflective structures in cephalopods, but only in Loliginid squids are they and the sub-wavelength photonic structures they control dynamically tunable, driving changes in skin color for camouflage and communication. The reflectins are block copolymers with repeated canonical domains interspersed with cationic linkers. Neurotransmitter-activated signal transduction culminates in catalytic phosphorylation of the tunable reflectins’ cationic linkers, with the resulting charge-neutralization overcoming Coulombic repulsion to progressively allow condensation and concommitant assembly to form multimeric spheres of tunable size. Structural transitions of reflectins A1 and A2 were analyzed by dynamic light scattering, transmission electron microscopy, solution small angle x-ray scattering, circular dichroism, atomic force microscopy, and fluorimetry. We analyzed the assembly behavior of phospho-mimetic, deletion, and other mutants in conjunction with pH-titration as an *in vitro* surrogate of phosphorylation to discover a predictive relationship between the extent of neutralization of the protein’s net charge density and the size of resulting multimeric protein assemblies of narrow polydispersity. Comparison of mutants shows this sensitivity to neutralization resides in the linkers and is spatially distributed along the protein. These results are consistent with the behavior of a charge-stabilized colloidal system, while imaging of large particles, and analysis of sequence composition, suggest that assembly may proceed through a transient liquid-liquid phase separated intermediate. These results offer insights into the basis of reflectin-based tunable biophotonics and open new paths for the design of new reflectin mutants with tunable properties.

## Introduction

Cephalopods such as squid and octopuses possess an optically dynamic epithelium, enabling complex camouflage and communication^1^^,^^2^. In addition to pigmentary chromatophores, these animals possess reflective cells – leucophores and iridocytes - that act as structural reflectors through the interaction of light with their sub-wavelength nanostructures^3^.While leucophores are broadband scatterers of white light, iridocytes reflect specifically colored, iridescent light by angle- and wavelength-dependent constructive interference from intracellular Bragg reflectors. The lamellae of these reflectors are densely filled with cationic block copolymer-like proteins called reflectins and are separated from the low refractive index extracellular fluid by regular invaginations of the cell membrane^4^^–^^8^.

Although the structural reflectors of the iridocytes and leucophores of most cephalopods are static, those in the Loliginid squid family uniquely possess reversibly tunable versions of these reflectors^9^. Ultrastructural characterization of unactivated tunable iridocytes showed their intracellular lamellae to contain a heterogeneity of discontinuous ~10-20 nm nanoparticles and nanofibrils, suggesting that the cationic reflectins within exist in a predominantly unassembled state dominated by inter-particle charge repulsion^9^^,^^10^. However, upon iridocyte activation initiated by binding of the neurotransmitter acetylcholine (ACh), released from nearby nerve cells, to cell surface muscarinic receptors, a signal transduction cascade culminates in enzymatic phosphorylation of the reflectins, consequently neutralizing their cationic charge and driving assembly of the reflectins to form homogenously densely staining Bragg lamellae^5^^,^^7^^,^^10^. Measurements demonstrating the reversible efflux of D_2_O and its re-uptake revealed that condensation of the reflectins drives the expulsion of H_2_O from the membrane-bounded lamellae, simultaneously increasing the refractive index contrast between the intracellular and extracellular layers of the Bragg reflector while shrinking their thickness and spacing, thus activating reflectance and progressively tuning the color of the reflected light across the visible spectrum^6^^,^^7^^,^^11^^,^^12^.

The role of the reflectins as a tunable driver of this biophotonic behavior has generated interest in understanding the principles underlying their responsiveness to signal-activated phosphorylation and their resulting changes in conformation and assembly. The reflectins are essentially block copolymers, being composed of highly conserved reflectin domains (RMs, for “Reflectin Motifs”) interspersed with cationic linkers, as seen for the reflectins A1 and A2 of the Loliginid squid *Doryteuthis opalescens* (**Figure 1A**). The reflectins exhibit a unique amino acid composition with some heterogeneity across their sequences, being highly enriched in methionine, arginine and tyrosine residues, and possess almost no (<1%) aliphatic residues^4^^,^^8^^,^^9^. The sequence composition of the reflectins suggests that attractive interactions are likely to be primarily driven by tyrosine-aromatic (pi-pi), arginine-tyrosine (cation-pi) and methionine-tyrosine (sulfur-pi) interactions, forming a complex interaction network of both intra- and inter-strand non-covalent bonding^9^^,^^13^.

**Figure 1.**
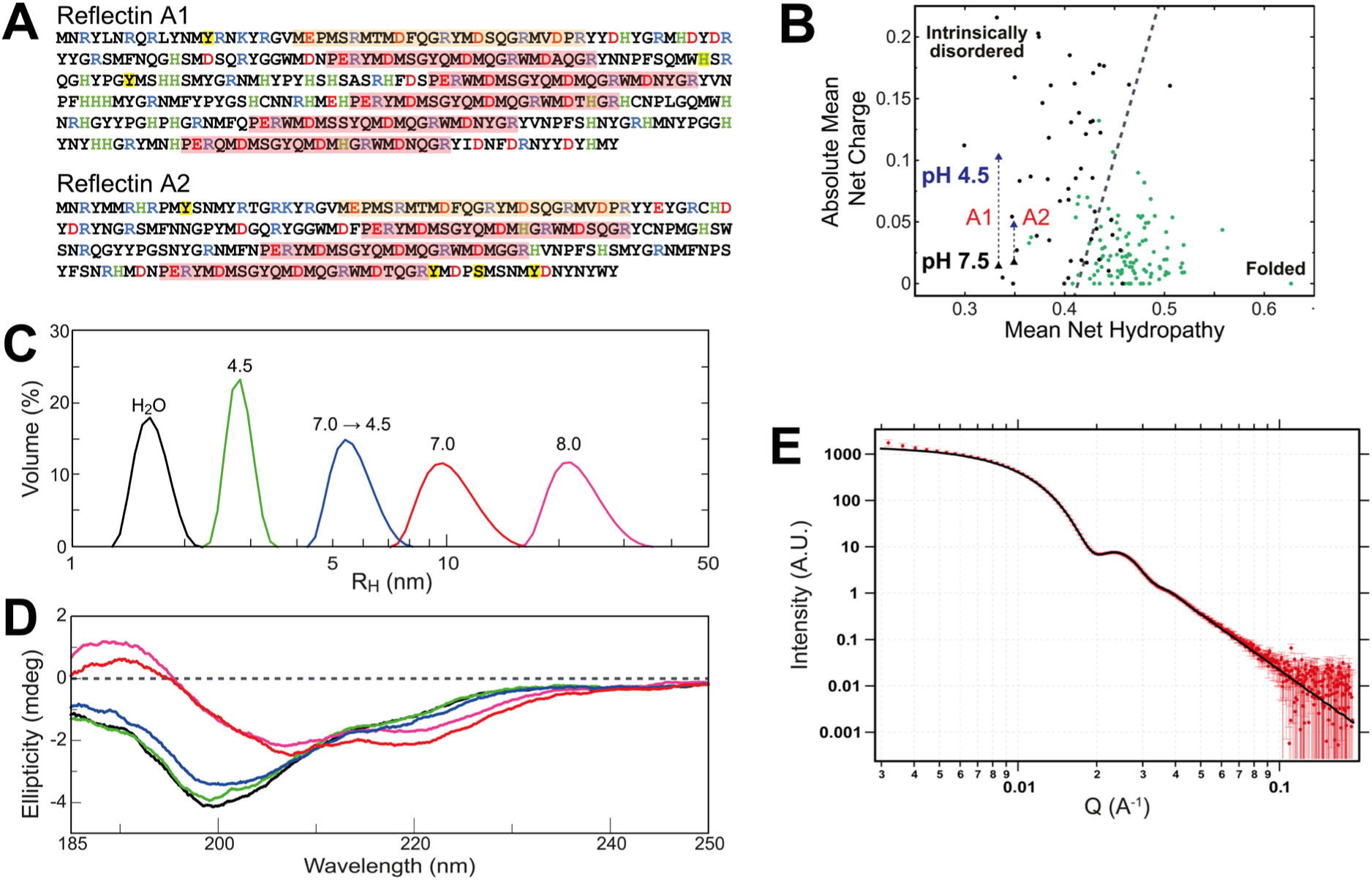
**A)** Sequences of *D. opalescens* reflectins A1 and A2. Conserved N-terminal domains are highlighted with dark yellow, other repeat domains with pale red. Histidine residues are green; other positively charged residues are blue; negatively-charged residues are bright red; phosphorylation sites are highlighted with bright yellow. **B)** Absolute mean net charge vs. hydropathy of a previously determined library of proteins (circles) and the reflectins (triangles). Yellow triangles designate mean net charge at neutral pH, where histidine residues are neutralized; red triangles designate acidic pH conditions, where histidines are presumed uniformly charged. In all other reference proteins histidines are considered neutral. Mean hydropathy calculated using the Kyte-Doolittle scale ^55^. **C)** Dynamic light scattering of A1 monomers (H2O and pH 4.5; black and green, respectively), multimers (pH 7.0; red), and oligomers (reversed from pH 7.0 → 4.5; blue). **D)** Circular dichroism spectra of A1 in monomeric, oligomeric, and multimeric states. Colors correspond to conditions as indicated for previous panel. **E**) SAXS data for reflectin A1 multimers (4 mg/mL) in 15 mM MOPS, pH 7.5, after subtraction of buffer background. Same sample was measured by DLS to have R_H_ = 20 nm. The data were fit to a spheroid model using SAXS modeling package IRENA, yielding an average particle radius of 23.2nm ± s.d. = 3.0nm.

Recently we demonstrated an *in vitro* assay that examined the assembly of purified recombinant *D. opalescens* reflectins as a function of progressive neutralization - as a surrogate of in vivo phosphorylation - through dilution of the H_2_O-solubilized proteins into low ionic strength buffers of varying pH^14^. Analyzing the tunable reflectins and a variety of mutant derivatives, we now show *in vitro* a predictive relationship between the extent of charge neutralization of the positively charged linker peptides and the size of the resulting assembled reflectin multimers. This discovery further elucidates the mechanistic origin of the synergistic effects of reflectin neutralization on the color and brightness of light reflected *in vivo.* Mutational analyses reveal that the “switch” controlling the neutralization-dependent structural transitions underlying tunability is not localized, but instead is spatially distributed in the multiple linkers along the reflectin’s length.

## Experimental Procedures

### Bioinformatics and Sequence Net Charge Calculations

Intrinsic disorder in the *D. opalescens* reflectins was predicted using the web-based servers FoldIndex^15^, PONDR-FIT^16^ and Metadisorder^17^ using default settings.

### Reflectin mutagenesis, expression, purification, solubilization, and buffer preparation

Recombinant *D. opalescens* reflectin A1 mutants were produced from a non-affinity tagged, codon optimized reflectin A1 WT construct described previously^6^^,^^14^. A1 mutants were generated by mutagenesis using standard techniques, and all mutants were confirmed by DNA sequencing. Codon-optimized DNA encoding the A2+5H/A2+10H mutants were purchased from ATUM (San Francisco, CA). Protein expression and purification were performed as described previously^14^. All proteins were expressed in Rosetta 2 (DE3) *E. coli* cells grown in half liter LB cultures from freshly plated transformants, in the presence of 50 mg/mL kanamycin and 37 mg/mL chloramphenicol. Expression was induced at A_600_ ~ 0.6, and allowed to proceed for ~6 h, at which point cells were pelleted by centrifugation and frozen at −80 °C until ready for use.

All reflectin proteins were expressed as inclusion bodies, similarly to WT, with roughly similar yields, as judged by comparison on SDS-PAGE gels (data not shown). Reflectin inclusion bodies were purified from cell pellets with BugBuster medium (Novagen, Inc., Madison, WI), as directed by the manufacturer. Inclusion bodies were solubilized in 5% acetic acic, 8M urea, 6M guanidinium-HCl, followed by dialysis against 5% acetic acid, 8M urea to remove guanidinium. Proteins were purified by ion exchange over a HiTrap XL (GE Healthcare) cation exchange column, and eluted with a gradient of 5% acetic acid, 6M Guanidinium. Fractions containing reflectin were collected, diluted in 5% acetic acid, 8M urea to lower guanidinium concentration, and then loaded onto a MonoS GL column (GE Healthcare), and eluted with a step gradient of 5% acetic acid, 6M guanidinium concentration. Eluted reflectin was concentrated and loaded onto a reverse phase C10 column by HPLC equilibrated with 0.1% trifluoracetic acid (TFA) in H_2_O, and eluted with a gradient of 95% acetonitrile, 0.1% TFA. Fractions were then lyophilized and stored in a −80 C freezer until solubilization. Purity was assessed on 10% Tris-acetate SDS-PAGE gels (Life Sciences, Carlsbad, CA).

### Protein solubilization and neutralization assay

Lyophilized protein was solubilized in 0.22 μm-filtered H_2_O. Protein concentration was determined by measuring absorbance at 280 nm using calculated extinction coefficients for each protein sequence. All freshly solubilized samples were diluted to 81 μM for use as stock solutions and stored at 4 °C between use. To perform the neutralization assay, all protein stocks, buffers, and water were filtered upon preparation and centrifuged at 18,000 x g for 10 minutes promptly before use and equilibrated to room temperature. Protein stock solution was diluted in prepared buffer solution to make final concentrations of 9-10 μM protein in 5 mM buffer. Sizes of reflectins are sensitive to buffer concentration, with higher buffer concentration leading to larger multimeric assemblies as a result of electrostatic charge screening; however, trends at higher buffer concentrations are consistent with those shown here. To determine reversibility of assembly, 250 mM acetate, pH 4.5 buffer was rapidly added and mixed to produce a final concentration of 15 mM acetate, pH 4.5.

### Dynamic light scattering

DLS analysis was performed on a Malvern Zetasizer Nano ZS (Worcestershire, Uk). All measurements were performed with pre-equilibrated samples at 25 °C. Multiple measurements per sample were taken to ensure sample equilibration and stability. All reflectin samples were measured at least three times, from at least two independent purifications, with variations generally less than 10% in assembly size, and results were reproducible across technical replicates and protein purifications. Data shown are representative.

### Circular Dichroism

All samples were measured on a Jasco J-1500 circular dichroism spectrophotometer using a 0.2 mm pathlength demountable cuvette (Starna Cells, Inc., Atascardero, CA). Reflectin monomer concentration in all experiments was 6 µM. Data shown are the averages of a minimum of 4 scans. For all data shown, high tension values were always less than 480 V.

### X-ray Scattering

Small- and Wide-Angle X-ray Scattering (SAXS and WAXS) measurements were conducted using a custom constructed instrument in the x-ray diffraction facility in the Materials Research Laboratory (MRL) at UCSB and at beamline 4-2 at the Stanford Synchrotron Radiation Lightsource (SSRL). The MRL instrument utilizes a 50 µm microfocus Cu target x-ray source (wavelength λ=0.154nm) with a parallel beam multilayer optics and monochromator (Genix, XENOCS, France), high efficiency scatterless hybrid slits collimator developed in house^18^, and a Eiger 1M solid state detectors (Dectris, Switzerland). Synchrotron SAXS data were collected at SSRL. Data reduction and model fitting were carried out using the NIKA and IRENA software packages developed at Argonne National Laboratory^19^^,^^20^.

### Transmission electron microscopy

All reflectin assemblies analyzed were prepared in 5 mM MOPS, pH 7.5 buffer, and measured by DLS before application to grids. 400-mesh carbon-coated grids (Electron Microscopy Sciences, Hatfield PA) were first treated by glow discharge for 30 seconds, and then 5 μL of freshly prepared sample was applied for 2 minutes before removal by wicking with filter paper. Samples were negatively stained 3X with 20 μL freshly-filtered 1.5% uranyl acetate for 15 sec followed by immediate wicking. Samples were visualized with a FEI Tecnai G2 Sphere Microscope at 200 kV. TEM images were processed in ImageJ 1.50i^21^.

### Atomic force microscopy

Assembled reflectin particles were applied for 30 sec onto cleaned glass slides before removal of buffer by wicking with filter paper. Negative controls with only buffer applied did not reveal any comparable particles. Reflectin particles were analyzed in tapping mode on a MFP-3D atomic force microscope (Asylum Research, Santa Barbara) with SSS-NCHR-10 high aspect ratio tips (Asylum Research, Santa Barbara). Data were processed using Gwyddion.

### Fluorescence

Samples were analyzed on a Cary Eclipse spectrophotometer (Varian, Palo Alto, USA), as described previously^14^. 200 μl of 9 μM samples were freshly prepared by neutralization assay before analysis. Excitation and emission slits were 5 nm width for all measurements.

## Results

### Bioinformatic analyses suggests the reflectins are intrinsically disordered

The tunable reflectins from the Loliginid squid *Doryteuthis opalescens* are essentially block-copolymeric, composed of unique and highly conserved peptide domains alternating with weakly polycationic linkers (**Figure 1A and Figure S1A)**^9^. Secondary structure algorithms do not predict significant alpha or beta structure, indicating that the reflectins are intrinsically disordered^4^^,^^5^. Comparison of *D. opalescens* reflectins A1 and A2 overall net charge and mean hydropathy with a collection of proteins previously analyzed for structure also indicates that the unmodified reflectins are intrinsically disordered (**Figure 1B**). Use of the FOLD-INDEX tool, which performs this same calculation using a moving window across the protein sequence, yields uniform scores across the entire reflectin A1 and A2 sequence, indicating that conserved domains and linkers do not vary in their drive to fold (**Figure S1A-B**). Use of the meta-predictors PONDR-FIT and Metadisorder support this current assessment, assigning either disordered or borderline ordered scores that show no correlatation with domain or linker sequences. (**Figure S1C-D**). These bioinformatic analyses agree with previous experimental analyses of reflectin nanoparticles by x-ray scattering, circular dichroism (CD), and Fourier-transform infrared spectroscopy (FTIR), as well as the CD analyses discussed below, all indicating that unmodified reflectins are largely disordered^6^^,^^22^^–^^25^. Interestingly, *D. opalescens* reflectins A1 and A2 show overall greater predicted levels of intrinsic disorder than reflectins from other cephalopod species, which do not display tunable iridescence (**Figure S1C-D**)^9^. This marginally greater drive of the *D. opalescens* reflectins towards disorder encoded within their sequences may play a role in enabling tunable assembly by shifting the equilibrium *in vivo* between the monomeric and assembled states, favoring disassembly in the absence of a triggering stimulus^9^.

### Unstructured monomeric reflectin reversibly forms relatively monodisperse nanoparticles

Dynamic light scattering (DLS) of 10 µM purified recombinant reflectin A1 in H_2_O-solubilized or acidic pH 4.5 conditions measures particles consistent with monomeric states (**Figure 1C**). Dilution of H_2_O-solubilized monomer into approximately pH 6.5+ conditions drives reversible assembly of spherical nanoparticle multimers of tunable size, estimated to contains between tens to thousands of reflectin monomers. This neutralization-driven assembly can be reversed by acidification with low concentrations of acetic acid, driving disassembly to form oligomeric particles with R_H_ values of approximately 6 nm, estimated to contain 10-17 monomers^9^. This estimate agrees with cryo-TEM analysis of an oligomeric non-tunable *Sepia offinalis* reflectin in the presence of SDS detergent^25^. Our DLS analysis of the *D. opalescens* A1 in 0.1% SDS confirms that they form stable monodisperse R_H_ ≈ 6 nm particles across our experimental pH range, showing that SDS stabilizes oligomers and prevents tunable assembly (data not shown). In the absence of SDS, these oligomers can be further cycled back to a multimeric state by dialysis into higher pH buffer, as shown previously. Circular dichroism (CD) spectra of monomeric *D. opalescens* A1 WT in both H_2_O-solubilized and acidic pH 4.5 conditions show strong minima near 200 nm and weak shoulders near 218 nm with both spectra, consistent with highly disordered states with a small degree of beta structure (**Figure 1D**). Multimeric assembly triggered by neutralization at pH 7.0 or pH 8.0 causes a substantial change in the CD spectrum, shifting the strong minimum and shoulder, suggesting significant structural changes during assembly. Notably, spectra of pH 7.0 and pH 8.0 multimers, which vary substantially in size (R_H_ = 10 nm vs 20 nm, respectively), display highly similar spectra, suggesting little pH-dependent conformational dependence. Acid-induced disassembly largely restores the CD spectrum to that of the disordered form, suggesting conformational similarities between oligomers and monomers.

### X-ray scattering of reflectin multimers

Synchrotron SAXS data collected from A1 samples in a multimeric state show well defined features characteristic of sphere-like particles (**Figure 1E**). The observed set of broad peaks in the scattering data can be attributed to form factor scattering arising from the Fourier Transform of a solid sphere. The data fit closely (solid line) to a model consisting of a group of randomly dispersed solid spheres, with an average radius = 23.2 ± 3.0 nm (approximately 13% polydispersity). The fit of the data to this model is remarkably faithful except at very low Q range, where the deviation can be explained by the presence of a small number of larger aggregates that were not included in the model. The average radius measured by SAXS agrees well with the R_H_ (20 nm) measured by DLS for the same sample. Samples prepared under different conditions exhibited a similar high degree of monodispersity, with calculated spherical radii also closely consistent with the R_H_ values measured by DLS (data not shown). Taken together, DLS, TEM, and SAXS provide a definitive confirmation that the multimeric particles are highly monodisperse spheres. Considering that the x-ray beam measures the ensemble average of the entire illuminated volume (~0.2mm x 0.2mm x 1.0mm) of the sample containing trillions of particles, the observed low size dispersion indicates the presence of a highly effective (and global) mechanism for size control of the multimer state.

### Design of reflectin mutants

Previous work showed that the tunability of *D. opalescens* reflectins is regulated by a neurotransmitter-triggered signal transduction cascade culminating in phosphorylation and consequent progressive charge-neutralization of the cationic reflectins, driving their progressive condensation and hierarchical assembly in a reversible and cyclable manner^5^^,^^7^^,^^9^^,^^14^. Plotting the distribution of electrostatics along the *D. opalescens* A1 sequence reveals a high degree of charge patterning across the reflectin linkers under both acidic and neutral conditions (i.e., conditions in which the histidines are either protonated or deprotonated, respectively) (**Figure 2A**). We find that these features are generally conserved across the reflectins found in diverse cephalopods (**Figure S2**). We generated and analyzed a collection of *D. opalescens* reflectin A1 mutants to better analyze the effect of charge on condensation and assembly *in vitro* (**Figure 2B**). A1 was chosen as the mutational platform based on its readily reproducible, tunably reversible assembly^14^. All mutants were expressed in bacteria and purified by methods identical to those used for the wild-type (WT), and no substantial differences from WT were observed during purification.

**Figure 2.**
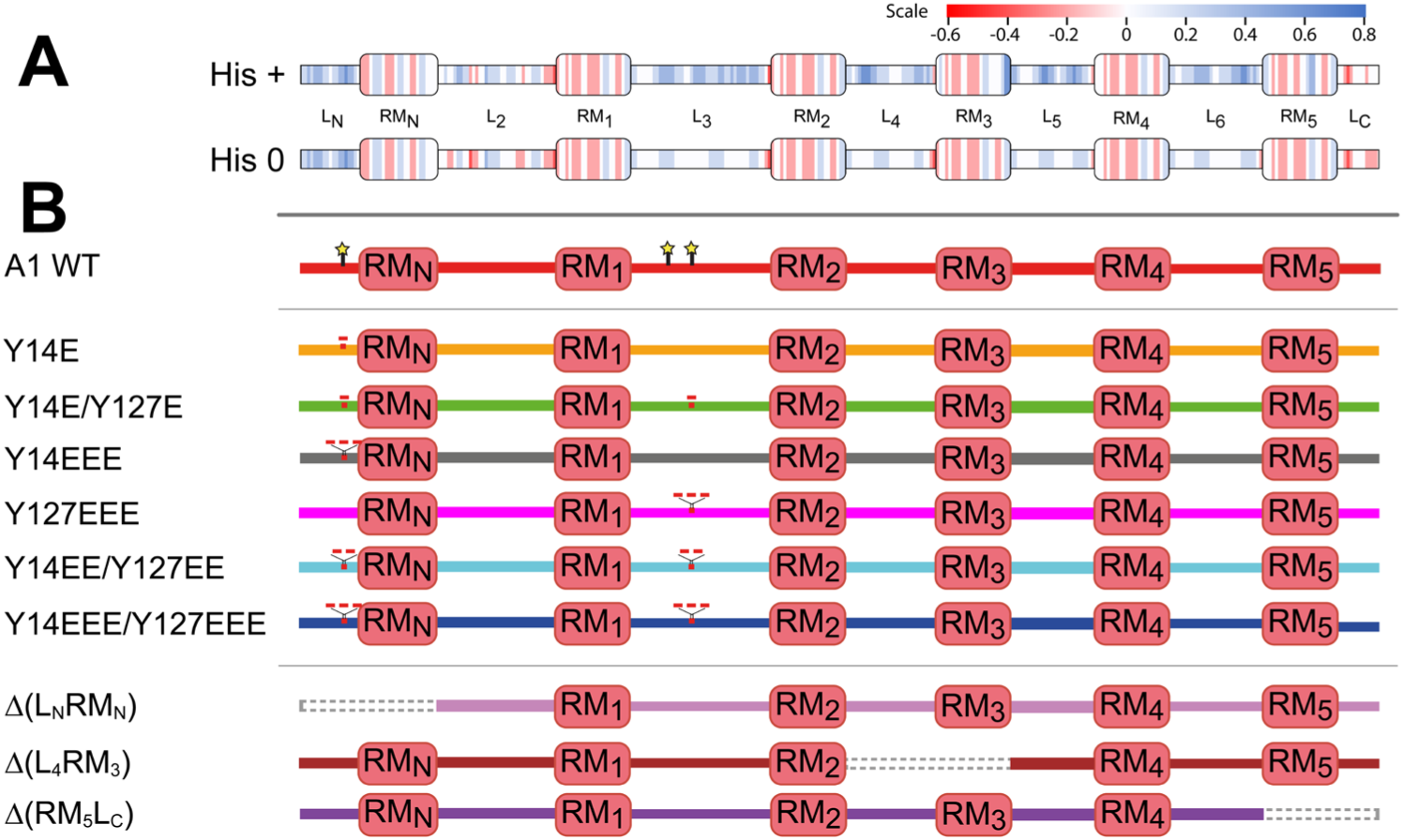
**A)** Scaled net charge across *D. opalescens* reflectin A1, in the histidine protonated (top) and histidine deprotonated (bottom) states. Color is plotted from a centered 5 residue moving window, using color key shown in figure. **B)** *D. opalescens* reflectin A1 mutationally altered proteins investigated in this work. Dotted gray lines designate deleted segments in respective mutants. In both **A** and **B**, boxes indicate conserved domains; lines indicate linkers.

Two classes of A1 mutants are investigated here. For the first, we created a progressive series of phosphomimetic glutamate mutations at two previously-identified *in vivo* phosphorylation sites (Y14 and Y127) found to be associated with the activation of tunable, reversible iridescence in dorsal *D. opalescens* iridocytes^8^. Additional numbers of added glutamate residues allowed investigation of the incremental effects of charge neutralization on assembly, as well as improved mimicry of the effects of the physiological addition of 2 negative charges with each covalently attached phosphate. The second class of mutants incorporated conserved domain-linker segment deletions, from either the N-terminal, central, or C-terminal sections of the protein. In *D. opalescens* A1, as in other reflectins (**Figure S2**), the N-terminal linker is highly positive charged, whereas the more central linkers exhibit a lower but significant net positive charge, and the C-terminus itself is negatively charged. To maintain the linker/motif balance seen in WT A-type reflectins, all mutants contained paired linker and conserved domain segment deletions. Due to the heterogenous charge distribution across reflectin A1, each conserved domain-linker deletion therefore uniquely modified the overall charge distribution of reflectin A1.

### Assembly behavior of reflectin mutants

We analyzed the pH-dependent assembly behavior of A1 WT and mutationally altered derivatives by DLS (**Figures 3A-H and S3&4**). Reliable interpretation of DLS R_H_ values requires particles to be spherical, a requirement that is justified by both the X-ray scattering discussed above as well as transmission electron microscopy (TEM) imaging of both WT and mutant reflectin assemblies, as discussed below. As judged by DLS (**Figure S3**) and TEM, as discussed below, both phosphomimetic (glutamate addition) and deletion mutants form largely monodisperse spherical assemblies morphologically similar to those previously observed for WT^14^. Also like WT, these assemblies formed rapidly, generally within seconds or less, although larger assembles with R_H_ > 100 nm showed growth by DLS for up to 20 minutes before stabilizing. Assemblies were stable over time (measured over several days), although R_H_ > approx. 100 nm particles settled and could not usually be recovered with resuspension. Sizes measured by DLS were reproducible, with standard deviations between assembly replicates generally < 10% of average measured DLS assembly size for a given condition when R_H_ ≤ 100 nm, although larger standard deviations between replicates (up to 49%) were observed for some large particle sizes (RH >> 100 nm).

**Figure 3.**
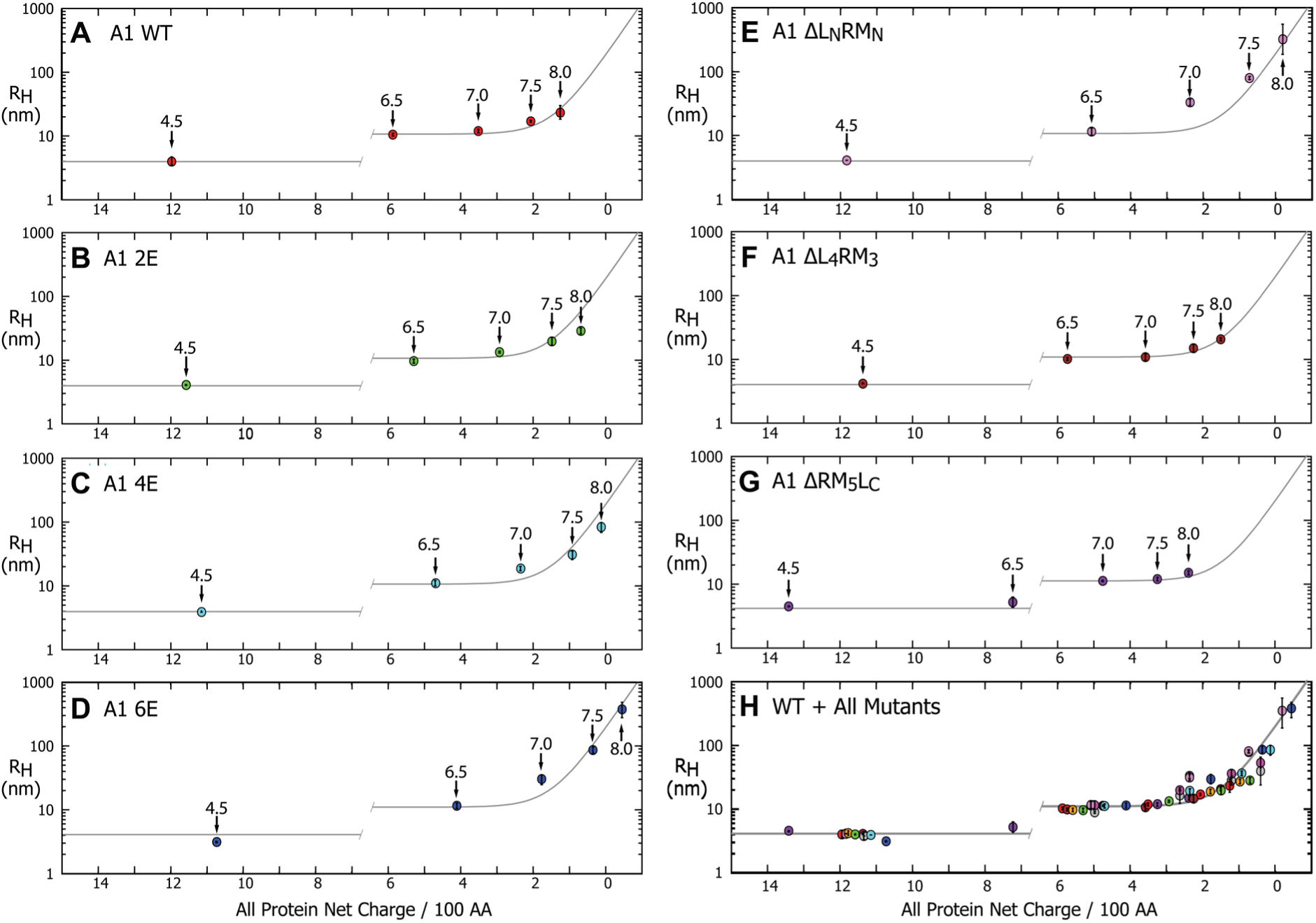
Experimentally determined R_H_ measured by DLS vs. calculated net charge density for: **A)** A1 WT. **B-D)** Selected A1 phosphomimetic mutants. **E-G)** A1 (conserved domain-linker)-deletion mutants. **H)** Data for WT and all mutants collected on a single graph. pH at each data point is labeled with arrows in panels. Discontinous gray line is the single exponential least squares fit of all A1 data for the multimers and plotted as flat for all monomers. Position of discontinuity between monomeric and multimeric states is estimated. Representative DLS intensity and volume data are shown in **Figure S3**; DLS size data against pH for all mutants are shown in **Figure S4**. Sizes are plotted as function of net charge density for each entire reflectin protein, as calculated from **Figures S5** and described in text. Each data point is the average of ≥ 3 replicate assembly measurements, where each assembly measurement is an average of ≥ 8 individual DLS measurements of a single sample. Error bars signify ± one standard deviation between technical replicates.

Significantly, reductions in the reflectin’s net charge through mutagenesis, mimicking the effects of *in vivo* neurotransmitter-activated phosphorylation *in vivo* resulted in large, systematic and highly reproducible effects on the size of assembly, with progressive neutralization reproducibly generating relatively monodisperse multimeric assemblies. Under identical pH conditions, mutationally altered reflectins with increasing numbers of glutamates formed progressively larger assemblies upon neutralization under identical conditions, with this effect being enhanced at progressively higher pH (**Figure 3A-D and S4A**). Both single-site single glutamate mutants, A1 Y14E and A1 Y127E, assembled to sizes in the range of A1 WT, showing that abolition of native tyrosines at those positions does not significantly perturb assembly. Further addition of negatively charged glutamates to A1 nonlinearly increased assembly size under identical pH conditions, with the effects of mutational neutralization producing exponentially greater effects as pH increases. Interestingly, both single-site triple glutamate mutants, A1 Y14EEE and A1 Y127EEE, had similar sizes across the entire range of pH tested, with these sizes being overall intermediate to A1 Y14E/Y127E (two glutamates added) and A1 Y14EE/Y127EE (four glutamates added), indicating a relative insensitivity to the precise localization of negative charge. A1 Y14EEE/Y127EEE, being maximally neutralized of these gluatamate mutants, resulted in the largest assemblies, with sizes up to R_H_ ≈ 350 nm, though it is worth noting that these assemblies were still relatively monodisperse and reversible by acidification by addition of 15 mM acetic acid, pH 4.5 – as were all glutamate mutants (data not shown).

Deletions of conserved domain-linker pair deletions also yielded reproducible effects on assembly size upon neutralization (**Figure 3E-G** and **S4B**). An N-terminal conserved domain-linker pair deletion (A1 ΔL_N_RM_N_) formed particles similar in size to the maximally-neutralized phosphomimetic A1 Y14EEE/Y127EEE, while middle and C-terminal conserved domain-linker pair deletions (A1 ΔL_4_RM_3_ and ΔRM_5_L_C_, respectively) assembled to slightly decreased sizes relative to WT. Notably, A1 ΔRM_5_LC formed R_H_ ≈ 4 nm particles at pH 6.5 of likely monomeric form, (**Figure 3G**), similar to pH 4.5 conditions for WT and other mutants, and unlike the multimers normally observed at this pH. Interestingly, deletion mutants showed inconsistant reversal upon acidification with 15 mM acetic acid, pH 4.5, with oligomers only being intermittently detectable by DLS (data not shown).

### Net charge density precisely determines reflectin assembly size

Strikingly, we found that the assembly behavior of the WT and all mutants from both classes fall on the same plot when graphed against the net charge density calculated for each protein. (**Figure 3**, particularly **H**). Our calculation of net charge density assumed invariant sidechain pK_a_’s for N-terminal, C-terminal, internal residues, and 6.5 for the histidine pK_a_^14^^,^^26^. Plots of calculated protein net charge and linear net charge density as a function of pH for A1 WT and the mutants discussed here are shown in **Figure S5**, each analyzed for the entire reflectin protein (**Figure S5A-D**), the linkers only (**Figure S5E-H**), and the conserved domains only (**Figure S5I-L**). An apparent discontinuity is observable between monomeric A1 at high net charge densities (typically acidic, acetate-buffered conditions), and the formation of multimers of tunable size. This suggests that a discrete neutralization threshold must be passed to trigger *D. opalescens* reflectin assembly, after which the predictive exponential relationship between net charge density and reflectin assembly size is revealed.

Decomposition of theses sequences into their component linker and domain segments shows that the net charge density of the conserved domains is poorly predictive of assembly size, while the net charge density of the linkers yields a predictive relationship with measured particle size analagous to that observed for the entire protein (**Figure 4A-C**). We thus conclude that the neutralization-sensitive “switch” resides in the linkers, which function like a sensor in this respect. As noted above, the observed sizes of the A1 mutants with 3 added glutamates were overall similar to each other and intermediate to mutants with 2 and 4 added glutamates, demonstrating an indifference to the location of neutralizing charges. Because reflectin assembly size appears insensitive to both the location and the means of charge-neutralization, whether through progressive glutamate addition or deletion or by pH titration, the switch therefore appears to be distributed among the spatially segregated linkers. These results, as illustrated in **Figures 3-5**, all are consistent with the behavior of a charge-stabilized colloidal system.

**Figure 4.**
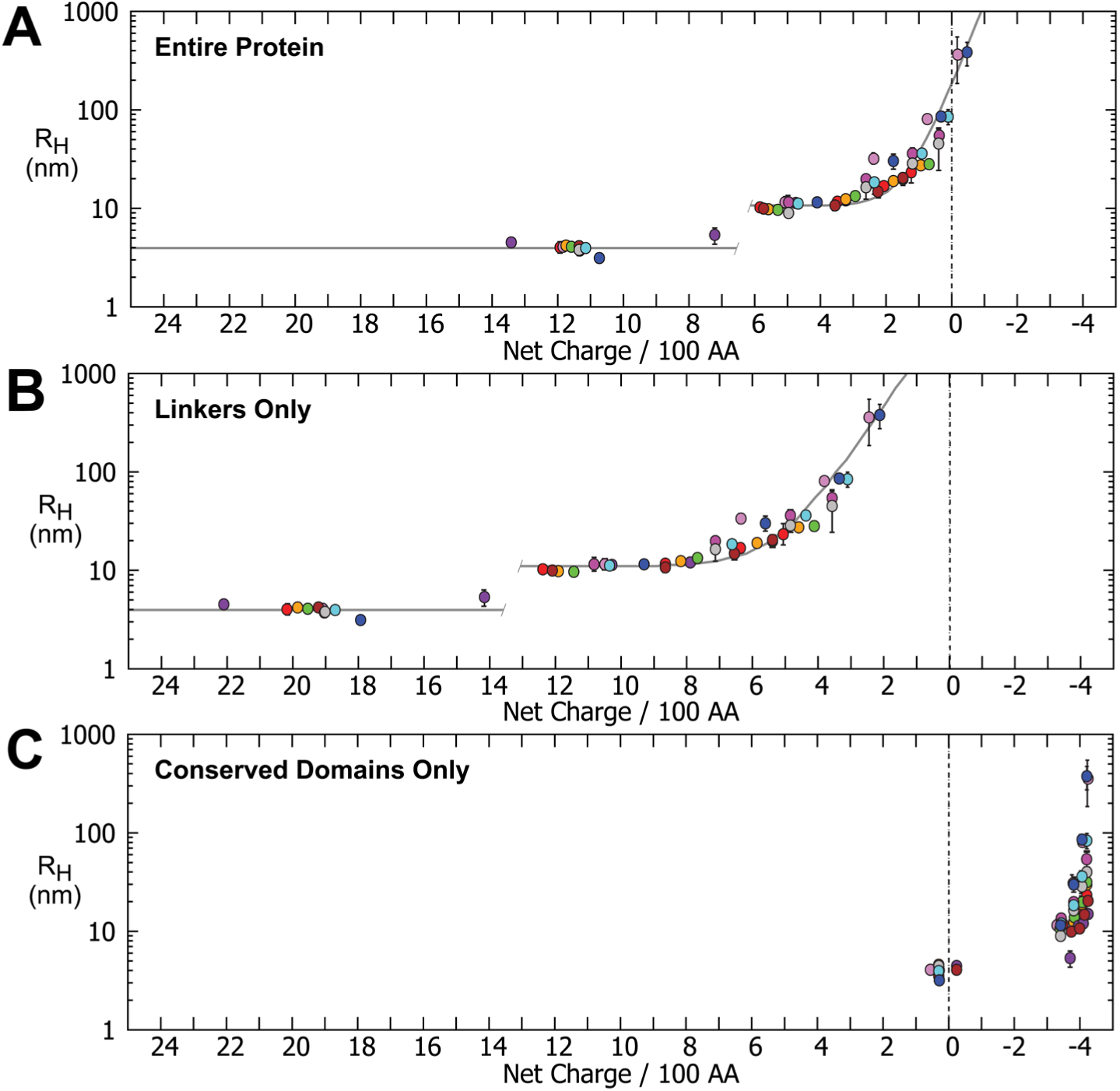
Experimentally measured R_H_ vs net charge density calculated for each entire protein or its conserved domains or linkers, for A1 WT and mutants at different pH values. Colors correspond to specific mutants as in **Figures 2 & 3**. Each data point is the average of ≥ 3 replicate assembly measurements, where each assembly measurement is an average of ≥ 8 individual DLS measurements of a sample. Error bars signify ± one standard deviation between technical replicates. Gray line is an exponential fit to all A1 multimer data, as in **Figure 2. A)** R_H_ as a function of net charge density calculated for entire A1 WT and mutant proteins. **B)** R_H_ as a function of net charge density calculated only for the linkers of A1 WT and mutants. **C)** R_H_ as a function of net charge density calculated only for the conserved domains (both RM_N_ and RM_#_s) of A1 WT and mutants.

### Characterization by electron microscopy, atomic force microscopy, and fluorescence

TEM analyses confirm that the reflectin A1 WT and mutants form assemblies of spheroidal morphology and relatively low polydispersity, with sizes measured by TEM agreeing well with those determined by DLS and SAXS (**Figure 5A-E)**. Similarity between A1 WT and A1 Y14E confirms that tyrosine mutagenesis itself does not generate aberrant assemblies. Particle size analysis of A1 WT and A1 Y14EEE / Y127EEE TEM micrographs indicate similar magnitudes of size variation (A1 WT polydispersity = 9%, A1 Y14EEE/Y127EEE polydispersity = 16%) (**Figure S6**). Imaging of the N-terminal conserved domain-linker pair deletion ΔLNRMN shows that it also forms spheres of a size consistent with the results from DLS. Close correspondence in assembly sizes measured in the hydrated state by DLS and upon drying on TEM grids suggests that the assemblies are stable and likely have low water content and internal dynamics, since they do not destructively shrink or collapse from drying upon sample application to the grid or exposure to vacuum in the electron microscope. Some larger assemblies, while appearing round in isolation, show distortions in morphology when packed into clusters, suggesting that they are deformable and do not coalesce (**Figure 5F**). Examination of air-dried reflectin particles by atomic force microscopy confirmed the highly symmetrical and smooth morphology of the solidified reflectin assemblies (**Figure 5G&H)**. Air-dried particles showed signs of flattening vertically with concommitent lateral expansion on the glass surface, demonstrating the same deformable nature observed by TEM for packed particles.

**Figure 5.**
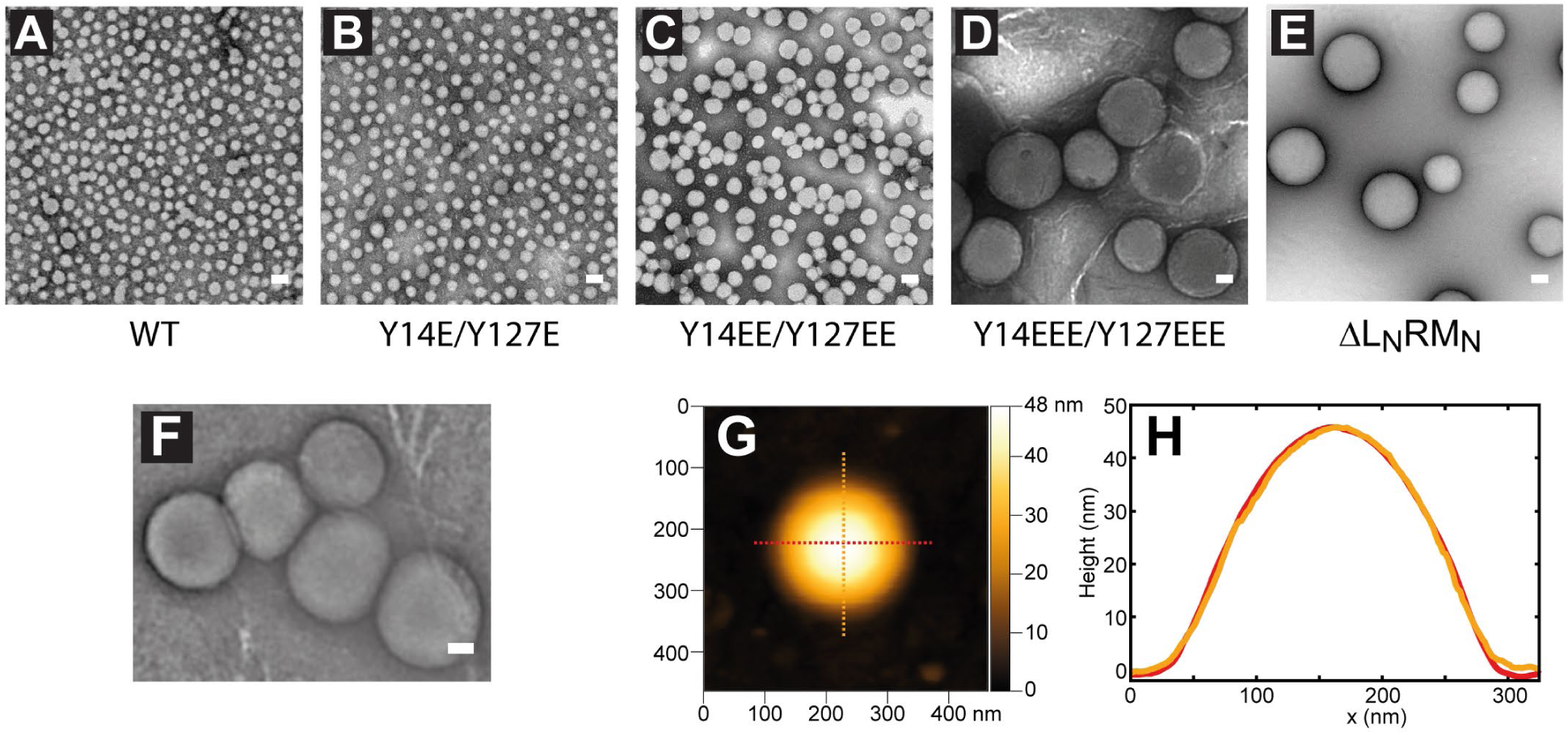
**A-E)** Transmission electron microscopy (TEM) images of A1 WT and glutamate mutants in 5 mM MOPS, pH 7.5. Scale bar = 50 nm. All TEM images in A-E have not been post-processed. Assemblies measured by DLS before application to the EM grids were: **A)** A1 WT; DLS R_H_ = 25 nm. **B)** A1 Y14E / Y127E; DLS R_H_ = 35 nm. **C)** A1 Y14EE / Y127EE; DLS R_H_ = 55 nm. **D)** A1 Y14EEE / Y127EEE; DLS R_H_ = 100 nm. **E)** A1 ΔL_N_RM_N_; DLS R_H_ = 77 nm. **F)** Cluster of A1 Y14EEE / Y127EEE particles, demonstrating deformation. Image was processed using a digital bandpass filter to enhance contrast^21^. **G)** Atomic force microscope image of an A1 ΔL_N_RM_N_ multimeric particle assembled in 5 mM MOPS, pH 7.5. **H)** Height profiles of portions of the A1 ΔL_N_RM_N_ particle shown in previous panel, as indicated.

The spheroidal and highly smooth appearance of the assemblies, particularly marked for larger assemblies such A1 Y14EEE/Y127EEE and ΔL_N_RM_N_, strongly suggests that a liquid-like phase transition occurs during assembly. However, the observed stability of these assemblies over time, their stability after drying, and the absence of any observed coalescence, would suggest that such a liquid-like phase possesses high interfacial energy and/or may rapidly vitrify in a reversible manner, although the deformable viscoelastic behavior of the large particles indicates that they are not entirely solidified.

*D. opalescens* reflectin A1 is strongly UV-fluorescent, as it contains 10 tryptophans, 7 of which are in highly conserved positions. Tryptophan fluorescence of the various phosphomimetic mutant nanoparticulate assemblies varied with charge neutralization-driven size in a manner essentially indistinguishable from that of WT (**Figure S7**; fluorescence of A1 WT was reported previously^14^), demonstrating that the substitution mutations produced no significant changes in internal structure observable by this method.

### Differences in net charge density explain differences between A1 and A2

Extending our application of net charge density beyond A1, we discovered that our previously-reported observation of differences between the assembly behaviors of reflectins A1 and A2 can now be understood. TEM shows that neutralized A2 assemblies have morphologies similar to those found for A1 (**Figure S8A**). Yet despite an identical pI and > 70% sequence identity with A1, A2 differed in response to addition of acid after assembly, with A2 being nonreversible^14^. Based on the experiments studying A1 assembly described above, we hypothesized that A2 assembly behavior can be readily understood to result from its lower percent histidine content; upon acidification, an insufficient number of histidines are protonated to sufficiently disrupt the assembly by Coulombic repulsion (**Figure 6A**). To test this hypothesis, we designed *D. opalescens* A2 mutants that contained an additional 5 (named A2+5H) and 10 (A2+10H) histidines spatially distributed among the linkers (**Figure 6A-C; Figure S8B**). We then compared the assembly and disassembly behavior of these mutationally altered reflectins with that of A2 WT as a function of pH (**Figure 6B&D**). Both histidine mutants of A2 assembled to similar sizes as WT, as expected based on net charge density, and followed the same net charge density curve derived from the A1 multimers (**Figure 6B**), supporting the conclusion that self-limitation of assembly size is determined by spatially distributed electrostatic interactions across the reflectin chain. However, histidine charge mutants differed significantly from WT in their disassembly upon acidification. While A2 WT was not reversible as measured by DLS in >90% of measurements, the A2+10H mutant protein was reversible in >90% of measurements (**Figure 6D&E**), forming R_H_≈12 nm oligomeric particles observable by DLS. Interesting, A2+5H showed variable behavior, being largely reversible at pH ≤ 7.0, and irreversible at pH ≥ 7.0, consistent with A2+5H possessing a net charge density near the charge threshold for assembly/disassembly.

**Figure 6.**
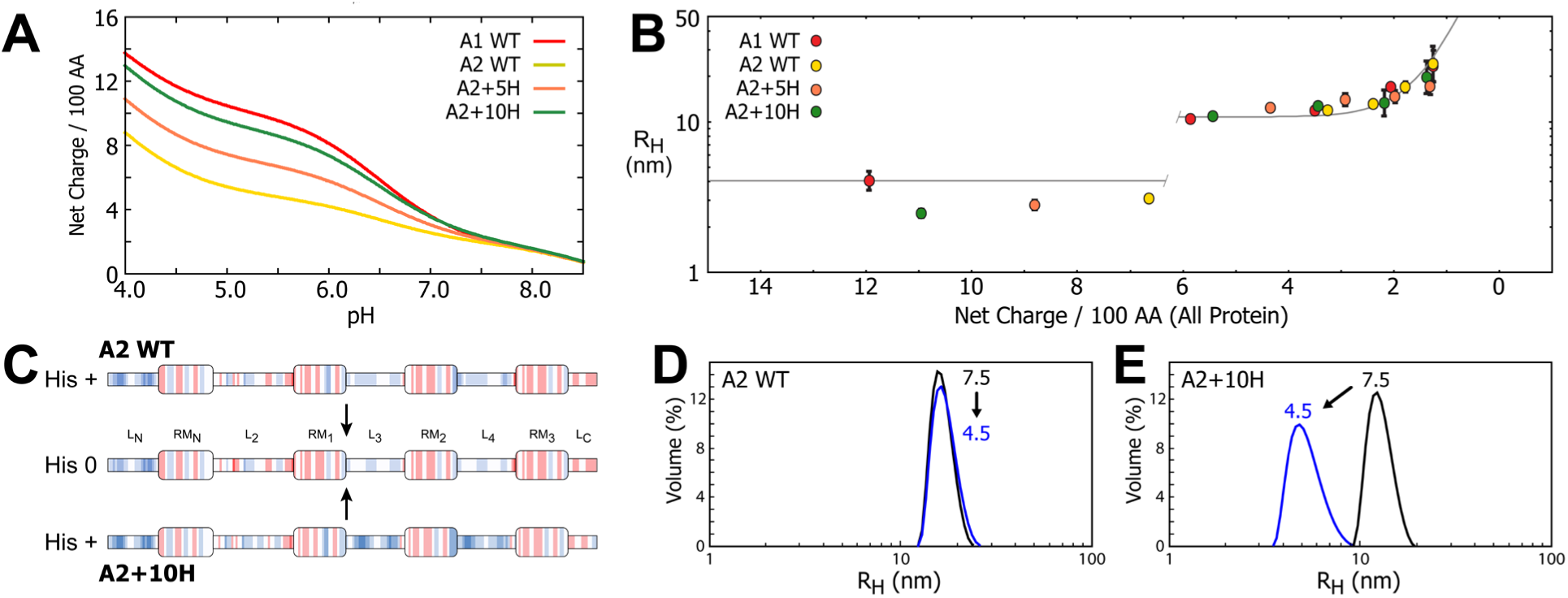
Comparison of *D. opalescens* reflectin A2 to A1 WT and mutants, and results for A2 histidine charge mutants. **A**) Net charge density calculated as a function of pH for A1 WT, A2 WT, and A2 mutants. **B)** Experimental R_H_ as a function of calculated net charge density of A1 WT, A2 WT, A2+5H, and A2+10H, under same pH range and conditions as A1 WT and mutants discussed earlier. Gray line is single exponential fit to all A1 multimer data, as in **Figures 2** **&** **3**. **C)** Comparison of charge distributions across A2 WT and A2+10H in histidine protonated and histidine uncharged states, as described for **Figure 2A**. Charge distributions of A2 WT, A2+5H, and A2+10H are indistinguishable in the neutralized state. **D-E)** DLS volume distributions showing non-reversibility of (**D**) A2 WT and reversibility of (**E**) A2+10H.

## Discussion

Reflectins assemble within cephalopod cells to form complex condensed nanostructures of high refractive index that produce diverse biophotonic effects^4^^–^^8^^,^^12^^,^^14^. In most cephalopods these structures are static, arising early during cellular development and remaining fixed for the lifetime of the cell^27^. However, within the Loliginid squids some iridocyte (iridescent / narrowband) and leucophore (broadband) cells demonstrate tunable, cyclable, acetylcholine-dependent reflectivity^5^^,^^28^. Ultrastructural and immunohistochemical characterization of iridocyte Bragg lamellae shows that unactivated tunable iridocytes are filled with a heterogenous network of small reflectin nanoparticles (ca. 10-20 nm diameter) and nanofibers^6^^,^^8^^,^^10^. Upon exposure to ACh, phosphorylation of reflectin A1 and A2 drives neutralization and concomitant condensation and assembly, resulting in expulsion of water from the membrane-bounded Bragg lamellae, thus increasing the intralamellar protein density while shrinking the thickness and spacing of the lamellae. These physical changes increase the refractive index constrast between the reflectin-containing lamellae and the interspersed extracellular medium to activate reflectance, while simultaneously actively tuning the reflected color^5^^,^^7^^,^^8^^,^^28^. Inspired by this unique biological function, studies have investigated the assembly and resulting optical behavior of purified recombinant reflectin and reflectin-based peptides. Purified reflectins and its peptides have been processed into a variety of optically active materials including thin films, gratings, and fibers^22^^,^^29^. These thin films have optical properties that are reversibly sensitive to both hydration and pH^22^^–^^24^^,^^29^^–^^32^. In a non-optical context, reflectin-based thin films have also been investigated as proton conductors and transistors, as well as substrates for neural cell growth^33^^–^^36^.

Starting from the highly disordered, H_2_O-solubilized reflectin monomers, we found that progressive neutralization drives the assembly of a collection of *D. opalescens* A1 WT and mutationally altered variants, with the resulting multimeric assembly sizes being predictively tuned by the normalized net charge density of the spatially distributed, cationic linkers that act as an electrostatic sensor and switch. Assembly size is indifferent to the mode of charge-neutralization (whether by genetic alteration, pH titration, or both – acting as surrogates of phosphorylation *in vivo*), and indifferent to the location of the cationic linkers remaining after deletion. TEM and SAXS analyses reveal the multimeric assemblies to be spherical and relatively monodisperse. The consistency of spherical morphology and internal structure (as measured from native tryptophan fluorescence) between WT and mutant reflectins indicates that the behavior of these mutant reflectins is not the result of aberrant or off-pathway aggregation, but lies on the common continuum, as also suggested by the colinearity of the data for the mutants and WT (**Figures 3&4**). Coulombic repulsion presumably contributes to maintenance of the positively charged reflectins in an extended, disordered and monomeric state, with progressive neutralization progressively overcoming that repulsion to permit condensation and assembly. The strong predictive relationship observed between the net charge density of the cationic linkers and reflectin’s final assembly size indicates that electrostatic interactions in some way limit that size.

To undergo assembly, the disordered reflectin monomers must pass through a net charge neutralization threshold, at which point they form multimers exhibiting an apparently exponential relationship between assembly size and calculated net charge density. Size control of reflectin assembly, tuned by changes in histidine protonation within the pH range tested here, emerges from a spatially distributed switch spread across the reflectin linkers. This behavior is consistent with the location of the previously identified sites of *in vivo* phosphorylation, almost exclusively found within the cationic linkers^5^^,^^8^. The reversible formation of oligomers in low pH / high net charge density conditions, either directly from an H2O-solubilized state or from larger multimer assemblies, suggests that assembly is hierarchical and that multimers may assemble from component oligomers. This would resemble the behavior of the intrinsically disordered amelogenin proteins, which have been observed to form monomers, oligomers, and larger nanoparticle multimers in a pH-controlled manner, and for which oligomers are thought to serve as the building blocks for larger nanoparticles^37^^,^^38^. However, unlike the amelogenins, the reflectins form monodisperse assemblies of tunable size as a function of neutralization, suggesting that some aspects of the unique reflectin amino acid composition and sequence enable a finer degree of control and tunability.

Comparison of *D. opalescens* reflectin A1 with similarly-sized representative reflectins from other cephalopods that have non-tunable iridescence shows that the N-terminal half of the reflectins generally possesses greater conservation (~50%) compared to the C-terminal half (~30%) (**Figure 7A)**. Interestingly, the N-terminal region, L_N_, possesses a well-conserved, strongly cationic character across all reflectins, being composed of ~19-26% arginine, compared to 11-12% arginine in the reflectins’ full sequences overall. The conserved cationic character of L_N_, combined with the experimental results reported here for the ΔL_N_RM_N_ mutant showing that it forms large assemblies, suggests that this linker may serve as an initial solubility tag during reflectin translation, inhibiting potential off-pathway aggregation. Our results show that linker electrostatics can broadly function to control reflectin condensation, consistent with the observed regulation *in vivo* regulation by phosphorylation. In these respects, the behavior of the reflectins is consistent with a charge-stabilized colloidal system, though additional complexity is apparent^39^^,^^40^.

**Figure 7.**
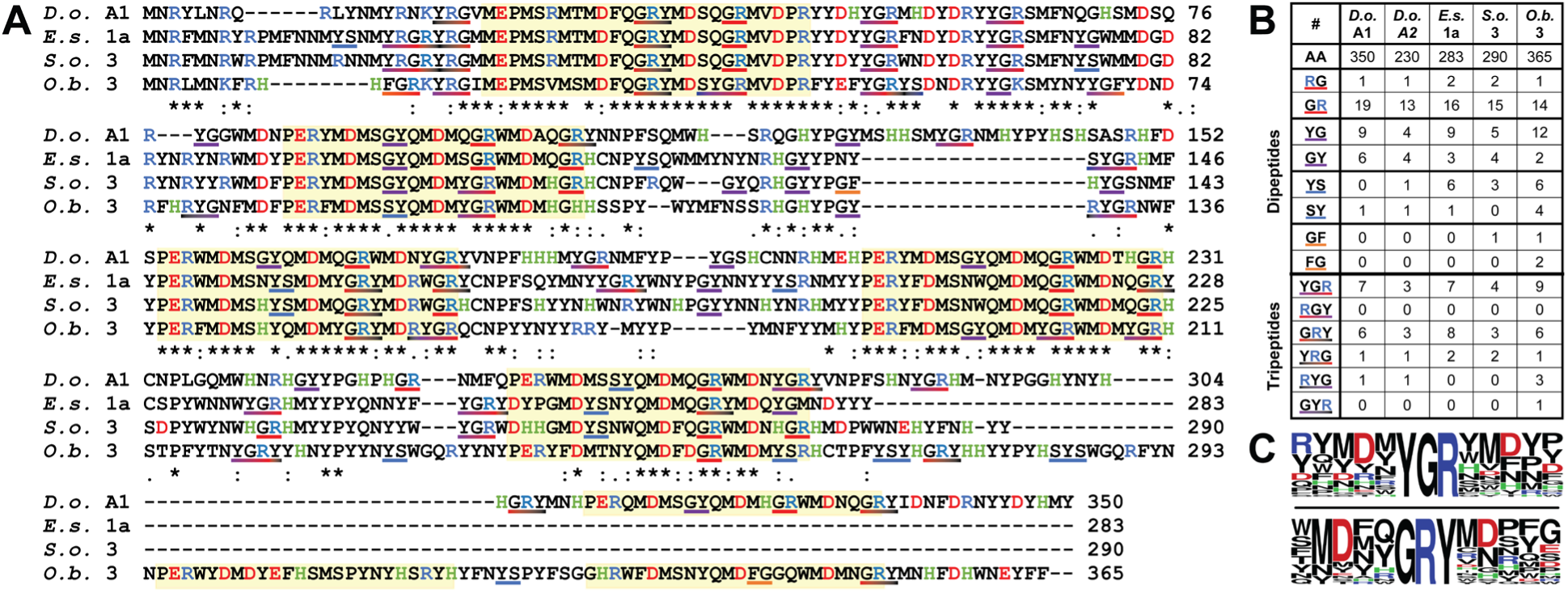
Analysis of di- and tripeptides associated with liquid-liquid phase segregation in representative reflectin proteins. **A**) Alignment of *D. opalescens* (squid) reflectin A1 (GenBank accession number KF661517.1), *Eupyrmna scolopes* (squid) reflectin 1a (AY294649.1), *Octopus bimaculoides* (octopus) reflectin 3 (KOF78298.1), and *Sepia offinalis* (cuttlefish) reflectin 3 (CCI88213.1). Conserved reflectin domains are highlighted in yellow. LLPS-associated di- (solid colors) and tripeptides (gradient colors) are underlined as labeled in table. Alignment performed using Clustal Omega^52^. **B)** Numbers of LLPS-associated di- and tripeptides in selected reflectin proteins. **C)** Sequence logos of regions surrounding YGR (upper) and GRY (lower) tripeptides seen in reflectins listed in **B**^56^.

Reflectins are highly enriched in conserved arginine (11%), tyrosine (20%) and methionine (15%) residues, and relatively deficient in lysines and residues with simple aliphatic sidechains (<1%). Thus, the reflectins are enriched in residues that are known to be associated with the liquid-liquid phase segregation (LLPS) of multivalent, intrinsically disordered proteins to form so-called biomolecular condensates, mediated through the formation of extensive webbed networks of relatively weak, shorter range cation-π, sulfur-π, and π-π associative interactions^41^^–^^43^. Arginine and tyrosine residues, often paired with glycine or serine, have been recognized as particularly strong drivers of protein LLPS^43^^–^^47^. An analysis of reflectin proteins from diverse cephalopods for di- and tripeptide sequence motifs associated with LLPS reveals a high occurrence of sequentially biased arg-gly, tyr-gly, and tyr-ser dipeptide sequences **(Figure 7A-B**), composing ~20% of the entire reflectin sequence^44^^,^^45^^,^^47^^,^^48^. Notably, these different dipeptide motifs are substantially clustered together to from a significant number of tyr-gly-arg and gly-arg-tyr tripeptides (**Figure 7A-C**). Given their high frequency and conservation in the reflectins, we suggest that these di- and tripeptides may play a critical role in driving reflectin LLPS. Based on the distribution of these associative peptide motifs throughout the reflectins, it is likely that both the conserved domains and the linkers robustly and directly participate in non-covalent bond formation during neutralization-triggered condensation.

While the high sphericity of the large reflectin A1 assemblies formed in vitro suggests that a transient phase-segregated liquid state may be formed upon sufficient charge neutralization and subsequence formation of reflectin condensates, the long temporal stability of reflectin assemblies and absence of coalescence and macroscopic phase separation as observed by DLS and light microscopy indicates that these particles do not progress to a state of low interfacial energy beyond a very short timescale, thus apparently undergoing phase separation arrest^14^^,^^42^^,^^49^. Solidification (or “vitrification”) of LLPS condensates is in several cases associated with disease states and is usually accompanied by deformations of the initially spherical assemblies, although a few exceptions are known^42^^,^^49^. For the reflectins, however, such non-spherical defomations may be avoided by a rapid solidification process. The charge-neutralization control of reflectin assembly sizes observed here may reflect the result of competition between constant attractive shorter-range attractive (e.g., cation-pi, electrostatic, etc.) interactions and tunable longer-range repulsive ones, which we experimentally tuned by histidine titration and manipulation of charges through mutation. This suggested mechanism is similar to one observed for some mussel foot proteins that are similarly rich in both cationic and aromatic residues, which have been shown to undergo LLPS upon an increase in ionic strength, postulated to act by screening the repulsive longer-range cationic interactions, while attractive shorter range cationic-pi interactions remain unaffected^50^. It remains unclear which aspects of the reflectin sequence and composition might enable the rapid solidification of condensates suggested here, although recent work with other proteins suggests that serine and glutamine content may be an important factor.^51^ The unusually high methionine content of the refectins suggests the possibility that a network of sulfur-pi bonds, acting in conjunction or parallel with the cation-pi network, may serve to sufficiently entangle reflectin monomers and slow internal dynamics.

It is likely that a mechanism of reflectin assembly and its regulation occurs within the Bragg lamellae of the tunable iridocytes that is analogous (although not necessarily identical) to that described here from our investigations of the purified recombinant proteins *in vitro.* The observation of apparently oligomeric nanoparticles and nanofibers in the unactivated tunable iridocytes, in comparison to the densely and homogenously staining proteins in the nontunable reflectin-based reflectors, indicates a pronounced difference in the degree of assembly of their reflectins: In the tunable iridocytes, the reflectins in the resting state appear oligomeric, while in the non-tunable iridocytes they appear multimeric^9^^,^^10^. While the exact mechanism underlying this difference remain unclear, it may result from an increased barrier to assembly in the tunable reflectins compared to the non-tunable reflectins, thus requiring phosphorylation (or otherwise neutralization *in vitro*) to induce assembly.

Biomolecular condensates have been found to play a wide variety of cellular roles, including signal transduction, cellular stress response, and RNA synthesis^42^^,^^43^. The tunable reflectins serve an apparently novel role as a molecular machine regulating an osmotic motor to drive the cyclable dehydration of Bragg lamellae, simultaneously increasing refractive index and shrinking the dimensions of the lamellae^7^. The reflectin’s unique sequence composition, rich in aromatic and sulfur-containing methionine, provides a further synergistic capability, providing the reflectins with one of the highest known incremental refractive indices (*dn*/*dc*), allowing the reflectins to generate higher refractive index contrast with the extracellular fluid in the physiological Bragg reflectors, while driving their tunability^9^^,^^52^.

The results described here highlight the reflectins as unusual and potential bioinspiration for future classes of tunably reconfigurable optical and other materials and may open further possibilities for exploration of the design principles underlying reflectin assembly; and suggest new pathways toward the rational design of tunable assembly^53^^,^^54^.

## Acknowledgements

This research was supported by grants from the U.S. Dept. of Energy, Basic Energy Sciences (DE-SC0015472), the Institute for Collaborative Biotechnologies through grant W911NF-09-0001 from the U.S. Army Research Office., and the Army Research Office (#W911NF-17-1-0160). The content of the information does not necessarily reflect the position or the policy of the Government, and no official endorsement should be inferred. We thank Cristophe A. Monnier for assistance with AFM measurements. The research carried out here made extensive use of shared experimental facilities of the Materials Research Laboratory, an NSF MRSEC, supported by NSF DMR 1720256. The MRL is a member of the NSF-supported Materials Research Facilities Network (http://www.mrfn.org).

## Author contributions

RL and DEM designed experiments and wrote the paper. RL, CB, CS, PK, YL, JS, and CA performed experiments. All authors reviewed results and approved the final version.

